# Widespread conservation of chromatin accessibility patterns and transcription factor binding in human and chimpanzee induced pluripotent stem cells

**DOI:** 10.1101/466631

**Authors:** Irene Gallego Romero, Shyam Gopalakrishnan, Yoav Gilad

## Abstract

Changes in gene regulation have been shown to contribute to phenotypic differences between closely related species, most notably in primates. It is likely that a subset of inter-species regulatory differences can be explained by changes in chromatin accessibility and transcription factor binding, yet there is a paucity of comparative data sets with which to investigate this. Using ATAC-seq, we profiled genome-wide chromatin accessibility in a matched set of 6 human and 6 chimpanzee (*Pan troglodytes*, our closest living relative) induced pluripotent stem cells from which we have previously collected gene expression data. We examined chromatin accessibility patterns near 20,745 orthologous transcriptions start sites and used a footprinting algorithm to predict transcription factor binding activity in each species. We found that the majority of chromatin accessibility patterns and transcription factor activity are conserved between these two closely related species. Interestingly, interspecies divergence in chromatin accessibility and transcription factor binding in pluripotent cells appear to contribute not to differences in the pluripotent state, but to downstream developmental processes. Put together, our findings suggest that the pluripotent state is extremely stable and potentially subject to stronger evolutionary constraint than other somatic tissues.

## Introduction

The contributions of gene regulatory differences to speciation and adaptation have been documented time and again by comparative genomics, aligning with the long-standing intuition that turnover at protein coding regions might not be sufficient to account for much of the phenotypic change among closely related species [1-5]. For example, ultra-conserved non-coding regions that exhibit accelerated change in a single lineage [6-9] have been shown to contribute to morphological differences between species [10-13]. In cases where the mechanism of action has been studied, the gain or loss of a single regulatory element can often be sufficient to drive phenotypes as substantial as the loss of limb development in snakes [14], or the loss of armour plates in fish [15].

Humans and other primates are no exception [5]. Human-specific gene expression and regulatory differences have been characterised during brain [16, 17] and limb development [18, 19]; likewise, steady-state differences in gene expression levels across myriad tissues have been repeatedly identified between humans and other primates [20-25] and attributed to changes in regulatory mechanisms such as histone modifications [26, 27], DNA methylation [28-30], repeat expansions [31, 32] (but see [33]), and differences in the activity of specific transcription factors (TF) [34]. A comparative study of DNaseI hypersensitive sites found more than 1000 loci that showed either human-specific or chimpanzee-specific gains or losses in hypersensitivity in fibroblasts and immortalised lymphoblastoid cells [35]. Many of these sites, which mark open chromatin presumed to be accessible to the cell’s transcriptional machinery, were located close to genes that also exhibited interspecies differences in their expression levels. Put together, a strong body of evidence suggests that regulatory changes and the associated differences in chromatin state make functional contributions to phenotypic differences between these species.

Two of the most intuitive mechanisms by which interspecies differences in gene expression levels can arise are through changes in chromatin accessibility and TF binding activity. Regulatory networks exhibit high degrees of robustness and resilience such that gene expression levels are generally conserved across species [36, 37], but multiple studies have shown that turnover at individual transcription factor binding sites (TFBS) is rapid, with few binding sites conserved across large evolutionary distances [38]. For example, a mouse model of human trisomy 21 carrying the entire human chromosome 21 alongside two copies of the mouse orthologue showed that expression levels of the genes on the human chromosome were more similar to those seen in human cells than to expression levels of their murine orthologues in the same cell. This finding argues strongly for a regulatory environment that is driven by DNA sequence rather than the nuclear environment [39]. Similarly, research in *Drosophila* has shown that changes in chromatin accessibility can be associated with changes in TF binding activity [40], and that chromatin accessibility and of TF binding sites can be used to predict gene expression divergence [41].

However, these previous studies have focused on only a handful of TFs, chosen because of their functional importance, rather than perform a general survey of the regulatory landscape. Direct high-throughput measurements of binding activity (using ChIP-seq, for example) across a representative suite of TFs in multiple samples – be they drawn from different tissues, individuals, or species – remain rare. In spite of recent technological advances, the need for validated antibodies, and/or large amounts of starting material often makes generating these data sets prohibitively expensive both in terms of labour and cost. An alternative popular approach, footprinting, combines chromatin accessibility data (originally in the form of DNaseI cleavage) with knowledge of a TF’s preferred binding sequence to predict whether a given site in the genome is bound or not [42].

Here, we profiled chromatin accessibility using ATAC-seq [43] in 6 human and 6 chimpanzee iPSC lines, which we had previously generated and extensively validated [44]. We first used these data to examine interspecies patterns of chromatin accessibility at 20,745 orthologous transcription start sites, and we examined the correlation of these data with existing gene expression data from the same cell lines. Then we applied a TF footprinting algorithm [45, 46] to 306 known position weight matrices (PWM), in order to perform an unprecedented characterisation of binding activity at over 130 million putative TF binding sites genome-wide. We found that the majority of chromatin accessibility and TF binding activity is conserved between these two closely related species, but in instances where it is not, interspecies change appears to contribute to downstream developmental processes.

## Results

To characterise the chromatin landscape in chimpanzee and human iPSCs, we generated ATAC-seq [43] libraries from six previously described lines from each species [44], and sequenced each sample to an average depth of ~252 million reads/sample. After exhaustive quality control and filtering of duplicate reads, mtDNA reads and reads outside high confidence orthologous regions (see methods), we retained an average of ~19.7 million reads per sample that mapped to a set of defined high-quality orthologous regions in the chimpanzee and human genomes (see methods and supplementary table 1). We additionally used the average ~99 million mtDNA reads from each sample to reconstruct their full mitochondrial genome sequences and infer the maternal lineage of our chimpanzee samples (see methods and supplementary figure 1), which was previously unknown.

### Changes in chromatin accessibility and gene expression levels

We initially focused our analyses on genomic regions that are expected to be accessible: transcription start sites. We began by defining 20,745 2kb windows centred on orthologous transcription start sites (orthologous TSS; see methods for more details). We also defined an equivalent set of control orthologous background genomic regions matched for broad mappability and located at least 5kb from any known TSS. Principal component analysis (PCA) of the orthologous TSS dataset does not show a clear visual separation between species (supplementary figure 2). While species is not significantly associated with any PC using the chromatin accessibility data from the orthologous background regions, only PC2 is significantly associated with species in the orthologous TSS dataset (PC2 *P* = 0.007; see supplementary table 2 for associations with additional PCs).

The results of the statistical analysis and the lack of clear visual separation of species in our data ran counter to previous observations of strong separation between species along the first PC when other types of regulatory data are used [44]. We thus sought an explanation. We compared the distribution of CPM values at orthologous TSS and orthologous background regions, and found that the former is clearly, and unexpectedly, bimodal (figure 1a; supplementary figure 3). By using k-means clustering on each species’ mean log_2_ CPM values, we found that of the 20,745 orthologous TSS in our dataset, 5,675 were assigned to a secondary CPM peak (figure 1b), which is associated with highly accessible regions. PCA using the data from only these 5,675 highly accessible TSS reveals a much stronger association with species than before (PC1 *P* = 4.0^∗^10^−5^; figure 1c), which can also be easily ascertained visually. The lack of clear separation of species in our overall data, therefore, has a technical explanation related to data quality (see methods for more details and additional relevant QC analysis), which impacts certain downstream analyses as we discuss below.

**Figure 1:**
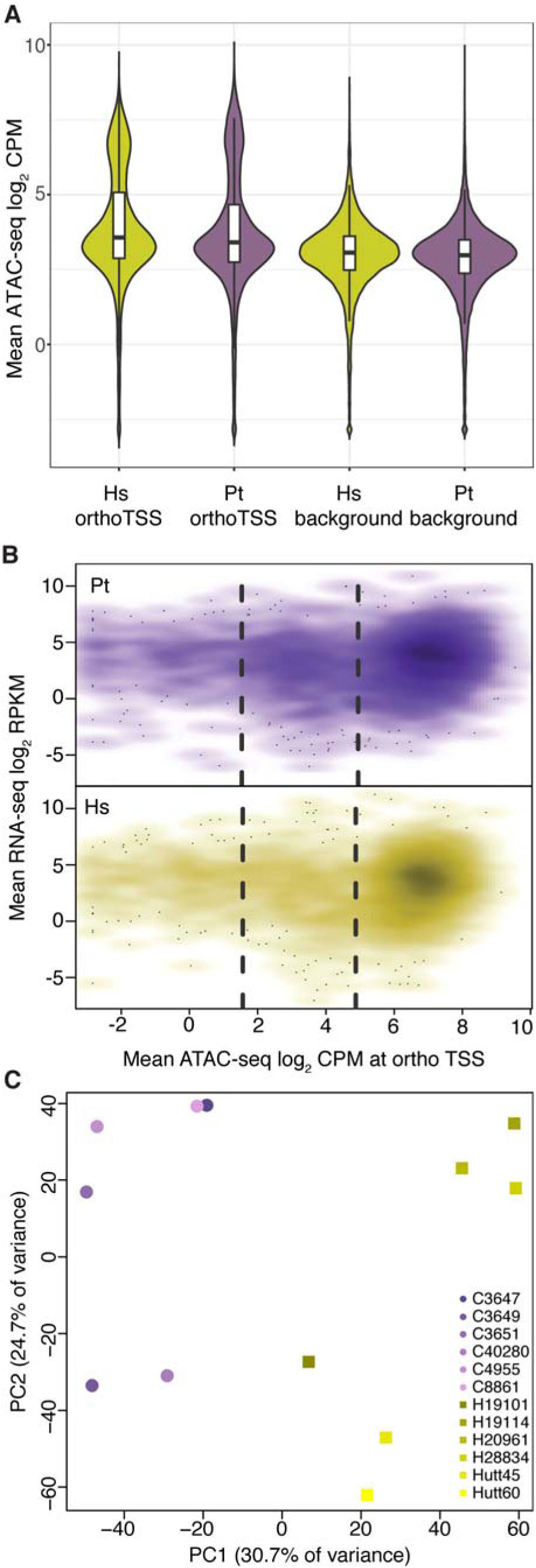
Patterns of chromatin accessibility in human and chimpanzee iPSCs. **a.** distribution of mean log2 CPM across 20,745 orthologous transcription start sites and 20,745 orthologous background regions. **b.** Expression (log2 RPKM) and accessibility (log2 CPM) in humans around the orthologous transcription start site of 4,210 genes in the highly accessible orthologous TSS subset expressed in chimpanzee (top) or human (bottom) iPSCs. The dashed lines indicate cluster boundaries identified through k-means clustering (k = 3) as described in the text. **c.** Principal component analysis of 5,675 orthologous transcription start sites.

To consider the ATAC-seq data alongside corresponding gene expression measurements, we re-analysed previously published RNA-sequencing data from the same cell lines [44], and mapped reads to an updated reference orthologous transcriptome (see methods and [21]). We detected the expression of 12,674 genes at log_2_ CPM >= 1 in at least half the individuals from one species, yet we were able to confidently identify an orthologous TSS for only 4,210 (33%) of these genes. This is due to our stringent definition of orthologous TSS, but the alternative is to introduce a strong bias towards regions with better mappability in humans than chimpanzees, which would impact our ability to detect genuine interspecies differences in chromatin accessibility. Of the expressed genes for which we can identify an orthologous TSS with confidence, 3,150 (75.0%) were genes whose orthologous TSS was part of the ‘highly accessible’ group defined above, a significant excess (hypergeometric *P* < 10^−16^) that supports this subset as being enriched for true biological signal (figure 1b).

Levels of accessibility at an orthologous TSS and the expression levels of the corresponding gene within an individual are weakly correlated (with Spearman’s ρ ranging between 0.05 for line Hutt60 to 0.17 for line H20961; *P* < 7^∗^10^−4^ for all individuals). The association improves somewhat if we only consider the 3,150 expressed genes with an orthologous TSS within the ‘highly accessible’ subset, but still remains modest: ρ ranges from 0.08 (line Hutt60; *P* = 2.6^∗^10^−6^) to 0.25 (line C8861G; *P* = 1.7^∗^10^−46^). This observation suggests that chromatin accessibility acts in a somewhat coarse fashion – remodelling is necessary to allow for transcription to occur, but the fine-tuning of expression levels likely occurs through additional mechanisms.

Bearing this in mind, we tested the 5,675 ‘highly accessible’ orthologous TSS (regardless of whether these TSS were associated with an expressed gene) for inter-species differences in accessibility, using an identical framework to that which we have previously used to detect differential gene expression [44]. At an FDR of 5%, 1,598 orthologous TSS are differentially accessible (DA) between the two species (supplementary table 3). Of these, 313 differentially accessible TSS are also associated with differentially expressed genes (supplementary table 4), a significant overlap (hypergeometric *P* = 8.6^∗^10^−3^, figure 2a). However, the relationship between inter-species differential expression and differential accessibility is not straightforward: Only in 64.5% of cases the inter-species direction of the effect is as expected, with the effect size sign being the same in in both the chromatin accessibility and gene expression data sets (figure 2b).

**Figure 2:**
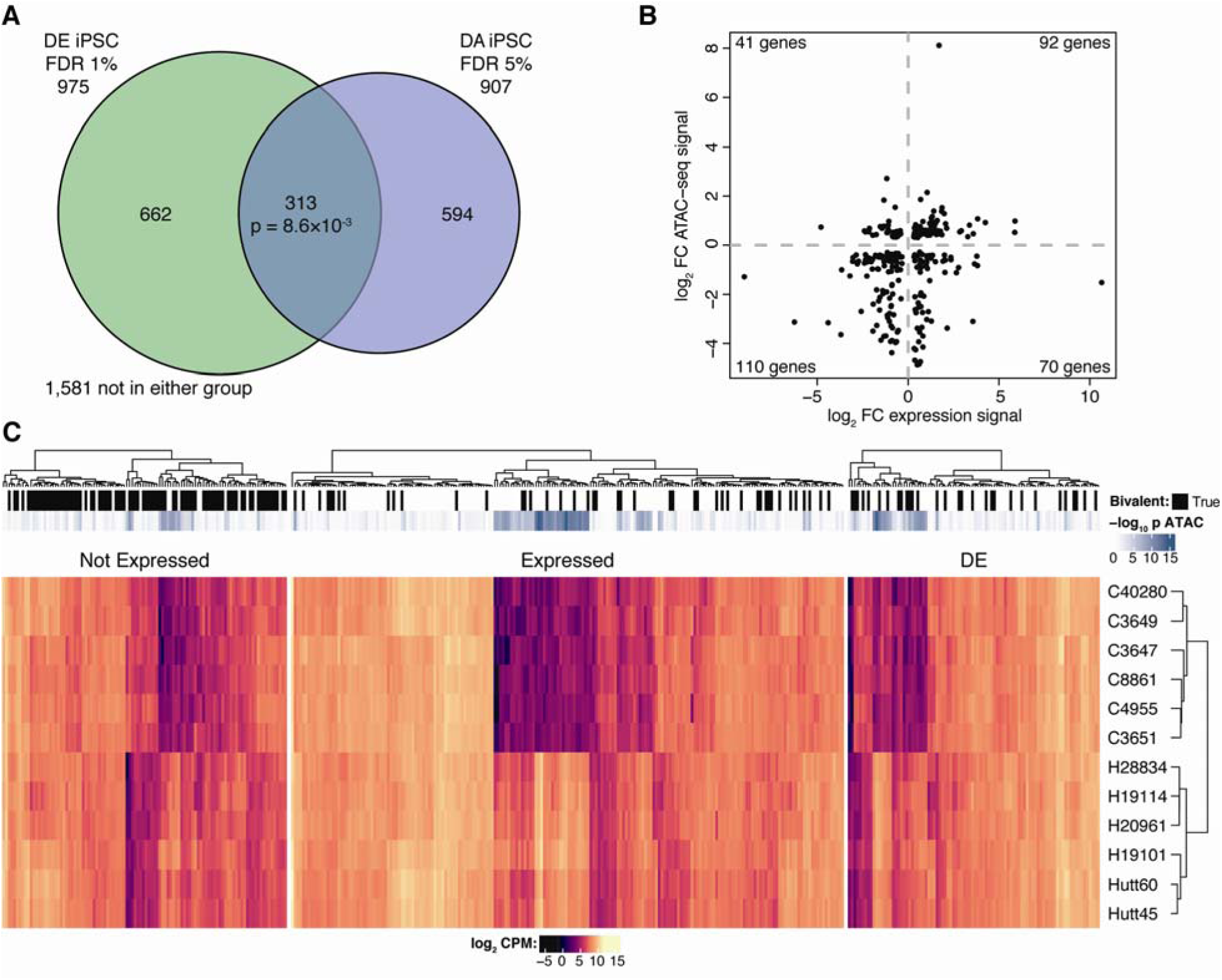
Testing for differentially accessible regions uncovers biologically meaningful signal. **a**. Overlap between differentially expressed and differentially accessible genes. p value from hypergeometric test. **b**. Effect directions across the 313 genes that are both differentially accessible and differentially expressed between human and chimpanzee iPSCs. c. Heatmap of 394 genes associated with significant GO terms, grouped by their expression status.

The 1,598 genes associated with orthologous TSS that are differentially accessible between species are enriched for five Gene Ontology Biological Process [47] terms, GO0007275: multicellular organismal development, GO0048731: system development, GO0044767: single-organism developmental process, GO0032502: developmental process and GO0060429: epithelium development (FDR = 10%). These enrichments are driven by a largely overlapping set of 398 genes, 294 of which are detectably expressed in at least one species, and 90 of which are differentially expressed between the two species (figure 2c; a full list is provided as supplementary table 5).

Interestingly, 104 of the genes associated with the orthologous TSS driving our GO results are not detected as expressed in iPSCs from either species. In order to understand the factors driving this observation, we asked whether these 104 regions might be playing a role in establishing gene expression patterns following exit from the pluripotent state [48]. To do so, we considered our data in combination with information on bivalent chromatin regions (“10_TssBiv”, “11_BivFlkn” and “12_EnhBiv”) identified in human embryonic stem cell line H1 as part of the Roadmap Epigenomics Project [49]. We found that 83 of these 104 orthologous TSS overlap at least one bivalent region in the Roadmap Epigenomics data, a far higher fraction than expected by chance alone (hypergeometric *P* < 10^−26^), as well as a high fraction compared to the overlap of bivalent regions with orthologous TSS associated with genes expressed in at least one species (75 out of 292; *P* = 0.97) or with TSS associated with differentially expressed genes (30 out of 90; *P* = 0.26). Indeed, the 104 genes associated with these TSS include multiple TFs with well-established roles in early development such as *ISL1*, which plays key roles in cardiac development [50], or *NEUROD2* and *NEUROD4*, both implicated in development of neural tissue [51-53], as well as other genes of less clear role, such as *RTN4RL1*.

#### Using ATAC-seq to infer inter-species differences in transcription factor binding activity

Because chromatin accessibility is not highly correlated with gene expression patterns either within or between species, we sought to characterise the landscape of TF binding activity in these cells by using footprinting analysis. To do so, we used a recent extension of the CENTIPEDE algorithm (msCentipede) that can better account for signal heterogeneity across sites (see methods; [46]), and predicted binding activity genome-wide across 306 position weight matrices (PWMs) from the HOCOMOCO database [54]. We considered 133,103,977 motif-predicted binding sites (MPBS) with a PWM score ≥ 7 in both the human (hg19) and chimpanzee (panTro2.1.3) genomes. This metric, which we calculated for each MPBS in each species, reflects how closely the locus matches the ideal PWM motif, with higher scores denoting higher fidelity. While each PWM is nominally associated with a single TF, there can be redundancy between PWMs associated with closely related TFs; thus we refer primarily to PWMs rather than TFs in the remaining sections.

We began by assessing the validity of our approach by comparing our binding predictions with all publicly released ENCODE [55] TF ChIP-seq data from the human embryonic stem cell line H1 (n = 28) and the human induced pluripotent stem cell line GM23338 (n = 5). Although recall and precision vary widely for different PWMs, we found that binding predictions from msCentipede often recapitulate findings for well-characterised PWMs and TFs (figure 3; additional details are available in supplementary table 6). For instance, ENCODE data contains three independent CTCF ChIP-seq experiments, one performed on H1 hESCs and two on GM23338 hiPSCs. On average, 25.1% of genome-wide CTCF MPBS in our data overlap ENCODE CTCF ChIP-seq peaks across at least one of the three experiments. When we considered only those sites msCentipede predicts to be bound in our human iPSCs, 73.7% overlap an ENCODE CTCF ChIP-seq peak, and 98.3% when we only considered bound MPBS with a PWM score ≥ 12 (a widely-accepted threshold for high quality MPBS). We found similar concordance when we considered MPBS for REST (also known as NRSF), NRF1, YY1 and GABPA amongst others.

**Figure 3:**
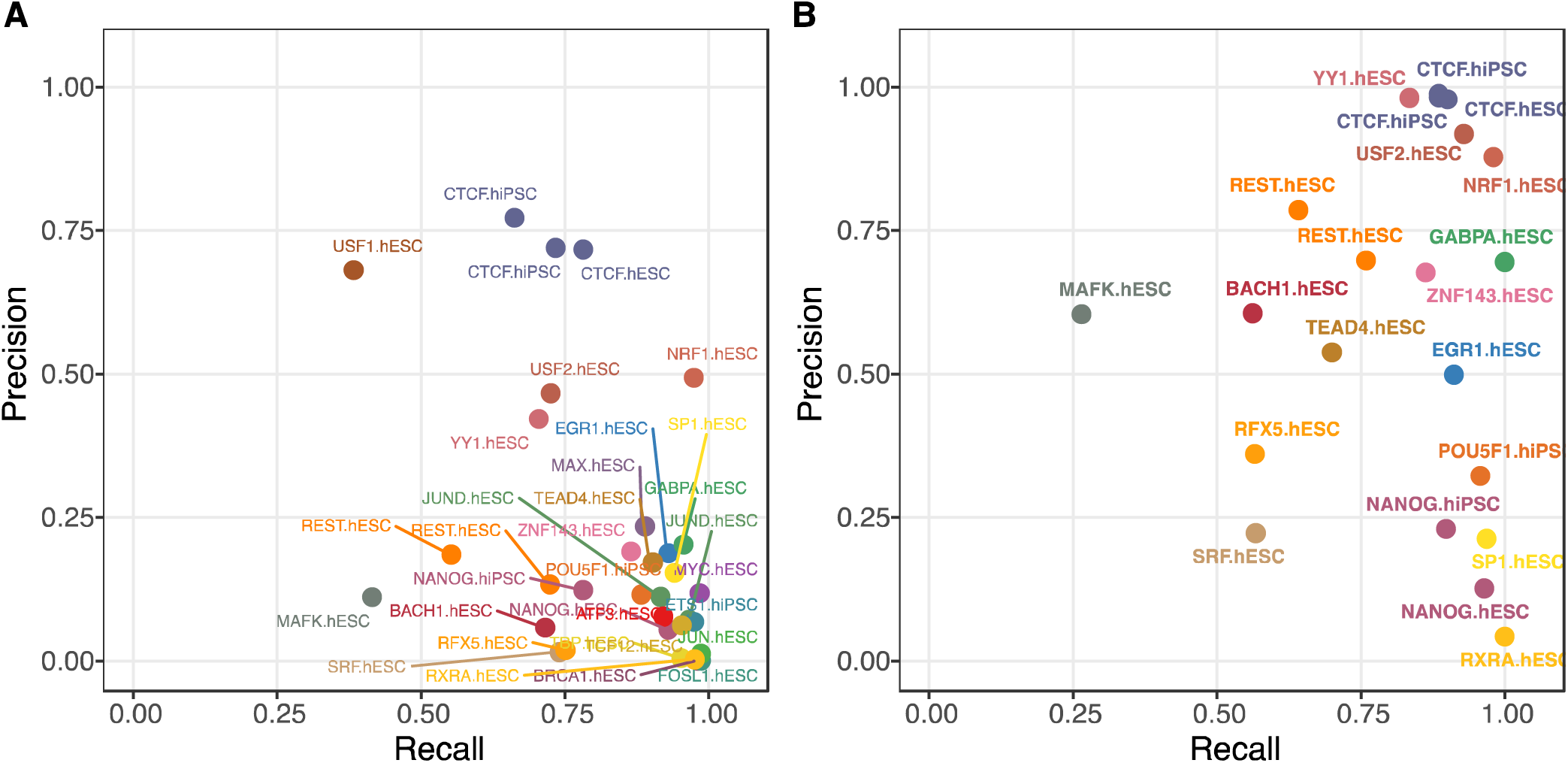
Recall and precision for msCentipede human binding predictions compared against ENCODE ChIP-seq data. **a**. At all sites with a PWM score ≥ 7. b. Within the subset of motifs with a PWM score ≥ 12.

Having validated our approach, we turned our attention to a comparison of MPBS across species. Across all PWMs, an average of 16.9% of sites are classified as bound (see methods) in humans, although values range from 0.7% to 97.3% for specific PWMs. In chimpanzee, an average of 9.5% of sites per PWM are classified as bound, ranging from 0.4% to 98.3%. The inter-species overall difference in binding can be explained by the lower read coverage in our chimpanzee samples, as discussed above (and in more detail in the methods). Reassuringly, an average of 85.5% of sites predicted to be bound in chimpanzees are also bound in humans (figure 4a). Thus we interpret the bulk of observed human-unique binding events as not being biologically meaningful but rather driven by technical differences, although we discuss some exceptions below.

**Figure 4:**
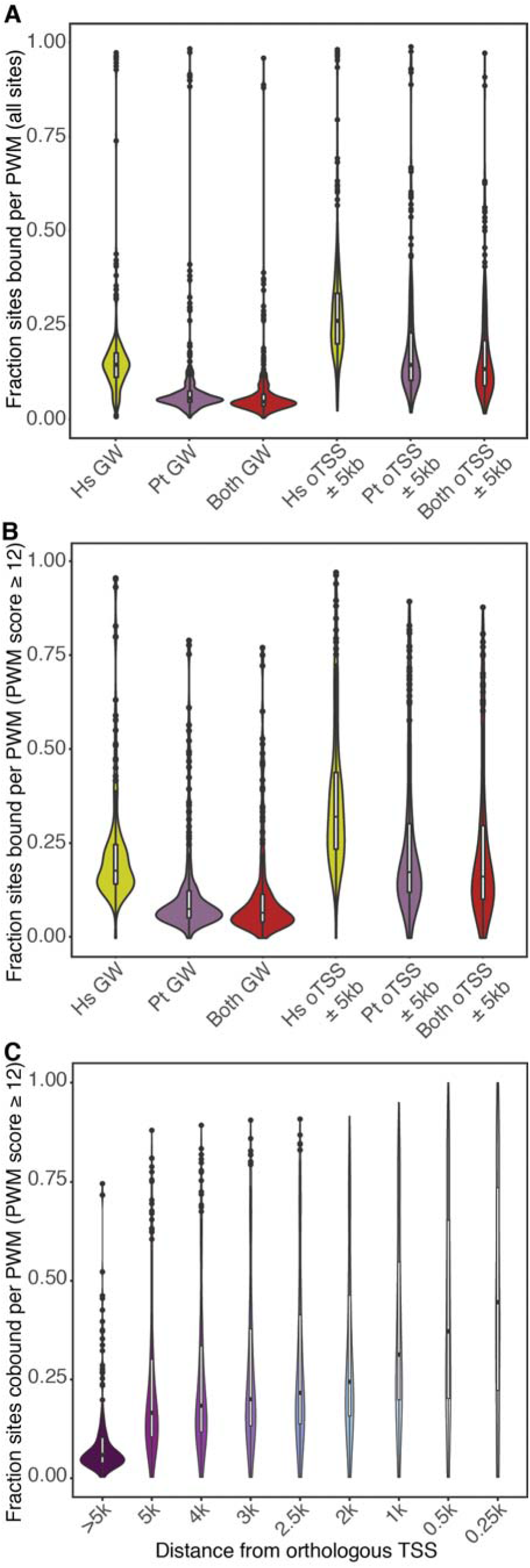
Transcription factor binding activity inferred by msCentipede in human and chimpanzee iPSCs. **a**. At all sites with a PWM score ≥ 7. **b**. Within the subset of motifs with a PWM score ≥ 12. **c**. Fraction of sites bound in both species at decreasing distance from orthologous TSS.

We limited our next analysis to the subset of motifs with a PWM score ≥ 12 (n = 906,612 across 173 PWMs). We found that, on average, 23.3% of sites are classified as bound in humans and 13.0% in chimpanzees. Of sites bound in chimpanzees, of 86.6% are also classified as bound in humans (figure 4b). We observed that predicted binding events do not occur randomly throughout the genome. When we classified binding sites by their distances to the nearest orthologous TSS, we found that binding significantly increases in frequency as distance to an orthologous TSS decreases (*P* < 1^∗^10^−16^; figure 4c). The concordance between human and chimpanzee results also increases for binding sites that are closer to the TSS. Interestingly, this observation is consistent across the vast majority of PWMs; the only exceptions are the PWMs associated with ERF and OTX2 in both species, PBX1 in humans only, and HINFP, TCF7L2 and NR3C2 in chimpanzees only (supplementary figure 4). Of these six TFs, *HINFP* and *TCF7L2* are differentially expressed between humans and chimpanzees; *ERF* is differentially accessible between the two species. However, all six PWMs are associated with small numbers of high quality MPBS (under 600 in all cases, as low as 32 in the case of HINFP), which may impact the predictive accuracy of msCentipede in these cases.

#### Unique binding patterns across species

There are multiple instances of predicted species-specific TF binding activity in our data. We explored possible mechanisms for divergence in TF binding by considering four scenarios: 1. *trans*-acting interspecies differences in the TF itself, possibly indicative of a change in motif preference, summarised by dN/dS values; 2. differences in the expression levels of the TF that binds the motif, also suggestive of *trans*-acting change; 3. differences in chromatin accessibility at MPBS, which can be tested by asking whether there is an excess of chimpanzee-unique binding events in regions differentially accessible between the species; and 4. *cis*-acting sequence turnover in the MPBS. To test this last possibility, we defined a simple metric, ΔPWM score, which is the difference at each MPBS between the PWM score in humans and chimpanzees.

Neither dN/dS ratios of TF divergence nor inter-species differences in the expression level of a TF are significantly associated with species-specific binding genome-wide. In contrast ΔPWM score between human and chimpanzee is associated with inter-species differences in TF binding, broadly suggesting that many predicted binding differences are due to *cis*-acting mechanisms. Sites inferred to be bound only in chimpanzees have, on average, a higher PWM score in chimpanzees than in humans, and vice versa (figure 5a). While the distribution of the ΔPWM score metric for sites bound in both species is centred at 0.002 (n = 146,554), the ΔPWM distribution for sites bound only in chimpanzee or only in human are clearly skewed (mean ΔPWM score for sites bound only in humans = 0.25; n = 107,784; *P* < 10^−16^; for sites bound only in chimpanzees = −0.29; n = 14,803; *P* < 10^−16^).

**Figure 5:**
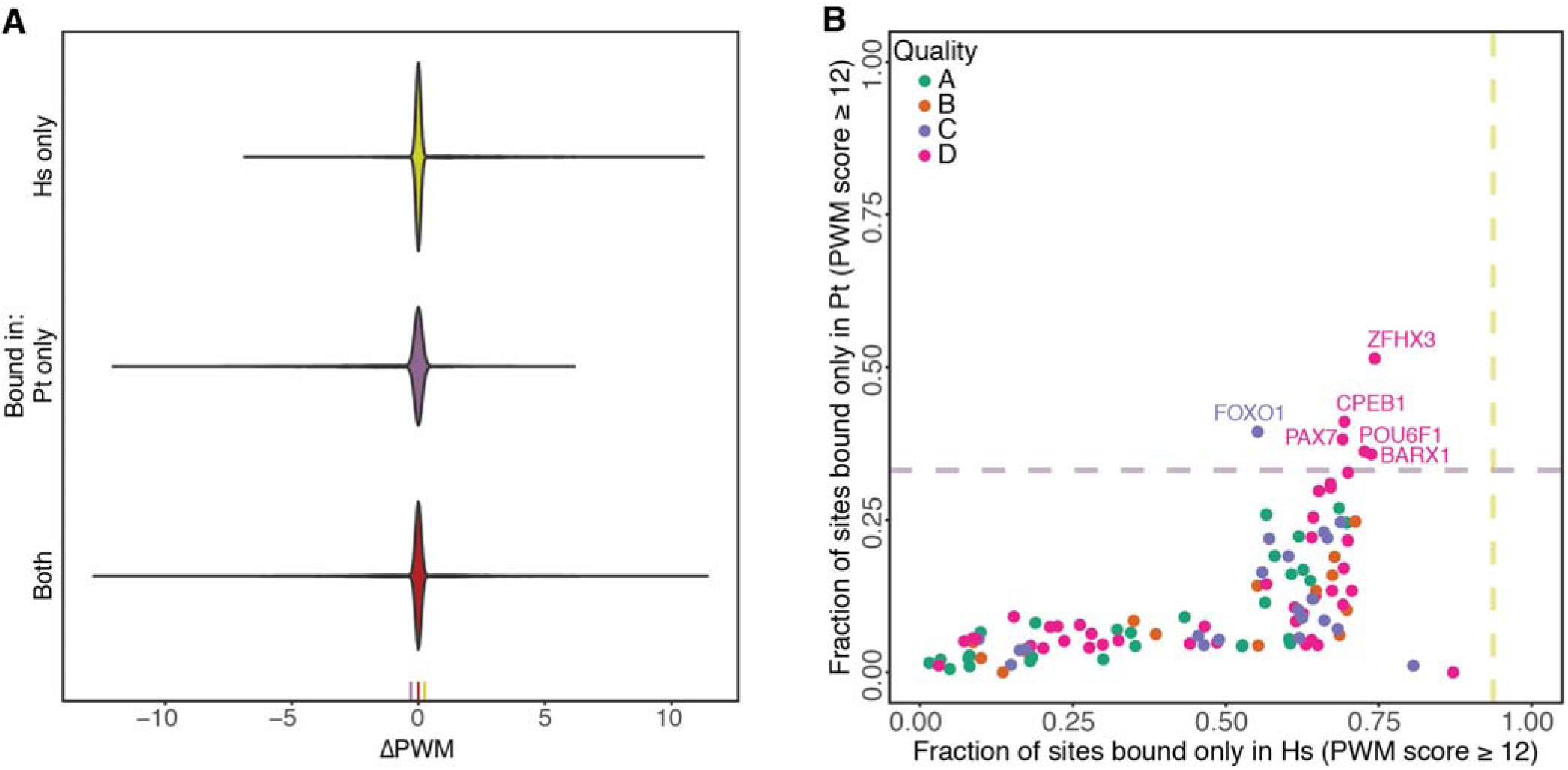
Binding differences between species. **a**. Distribution of ΔPWM scores by binding status. The three marks at the bottom highlight the median value for each dataset. **b**. Species-unique binding by PWM. The dashed purple and yellow lines denote 2 standard deviations away from the species-wide average for chimpanzee and humans, respectively.

We then focused on chimpanzee-specific binding events, as we expect these to more likely be true positive unique events (in contrast to the human-specific events, many of which are likely to be missing from chimpanzee due to technical reasons, as previously discussed). We asked whether chimpanzee-specific binding events around orthologous TSSs can be associated with differences in corresponding gene expression levels or chromatin accessibility. We considered all orthologous TSS (± 5kb) with a least one predicted high-quality chimpanzee-binding event (but not necessarily chimpanzee-specific) and with matched accessibility (n = 3,870) or expression (n = 2,981) data. We found a weak but significant correlation (Pearson’s R = 0.06; *P* = 2.0^∗^10^−4^) between interspecies differences in accessibility around orthologous TSS and the number of chimpanzee-specific binding events in the region. The magnitude of these differences is small: the mean number of chimpanzee-specific binding events for orthologous TSS that are not differentially accessible (DA) is 0.57 (n = 3,326), 0.83 for all DA orthologous TSS (n = 544), and 1.42 for DA orthologous TSS with an absolute log_2_ fold change in accessibility ≥ 2 (n = 19). We also found a weak correlation (Pearson’s R = 0.05, *P* = 0.01) between interspecies gene expression effect size (absolute log_2_ fold change) and the number of chimpanzee-specific binding events occurring within 5kb of the corresponding orthologous TSS. Again, the size of this effect is small: the mean number of chimpanzee-specific binding events for non-DE genes is 0.44 (n = 2,043), 0.53 for all DE genes (n = 938), and 1.03 for DE genes with an absolute log_2_ fold change ≥ 2 (n = 62).

We asked whether any PWM is associated with a systematic excess or deficit of species-specific binding genome-wide, which could suggest broad regulatory network rewiring and turnover. Again we focused on chimpanzee-specific excesses, and considered 109 PWMs with at least 100 predicted binding events in chimpanzees at high quality MPBS. We found 6 PWMs that exhibit chimpanzee-specific binding at levels more than 2 standard deviations away from the mean, associated with the TFs BARX1, CPEB1, FOXO1, PAX7, POU6F1 and ZFHX3 (figure 5b). These PWMs are ranked as class D by HOCOMOCO, suggestive of poor predictive value, with the exception of FOXO1, which is class C (see methods for more details). We did not find any PWMs that exhibit excessive human-specific binding. However, if we simply rank PWMs by their amount of human-specific binding we found that, with the exception of FOXO1, the same PWMs as above are amongst those with the greatest amount of specific binding in humans (figure 5b) potentially suggesting that these PWM generally evolve rapidly in the two species, or, alternatively, that in these cases the PWM does not capture the full scope of binding activity.

Of the 6 TFs associated with these PWMs, 3 are differentially expressed between human and chimpanzee iPSCs – *FOXO1*, *BARX1* and *PAX7*, although we found no excess of differentially expressed genes relative to the dataset-wide average (permutation *P* = 0.7). This observation might be explained by high redundancy in the binding motifs. For example, 399 of 1,014 binding events associated with the FOXO1 PWM in chimpanzee iPSCs are unique to the species. But the PWM itself has a high similarity to other sites associated with different forkhead family members such as *FOXJ3*, which is expressed in both humans and chimpanzees at high levels and with which *FOXO1* shares ~30% of possible high-quality binding sites.

#### Dynamics of activity in the core pluripotency regulatory network

Finally, we focused on PWMs associated with TFs with key roles in maintaining the pluripotent state. The TFs *OCT4* (also known as *POU5F1*), *SOX2* and *NANOG* sit atop the gene regulatory network of pluripotency, and act cooperatively to maintain this state [56]. Although both our previous work and that of others [44, 57] has shown that the pluripotency gene regulatory network is highly conserved between humans and chimpanzees, we found that conservation in the binding locations of these PWMs is not particularly high. Only 57.9% of OCT4, 55.1% of SOX2 and 55.2% of NANOG sites in the high-quality motif subset predicted to be bound in chimpanzee are also predicted to be bound in human. These observations, however, must be tempered by the poor concordance between the sites msCentipede predicts to be bound and ChIP-seq peaks for OCT4 and NANOG (both profiled in GM23338 iPSCs) in ENCODE data (figure 3), which in turn are likely driven by the complex binding dynamics of these TFs [58].

We also considered a larger set of 15 master pluripotency regulators drawn from the literature (specifically, from [56, 59, 60]). In this case, we found only few instances of chimpanzee-specific binding, suggesting overall high conservation of pluripotency pathways between the two species (mean fraction of chimpanzee-specific binding = 4.6% at high-quality sites, 4.9% across all sites; figure 6). This result is consistent with our previous observation that master pluripotency regulators are generally not differentially expressed between iPSCs from the two species [44].

**Figure 6:**
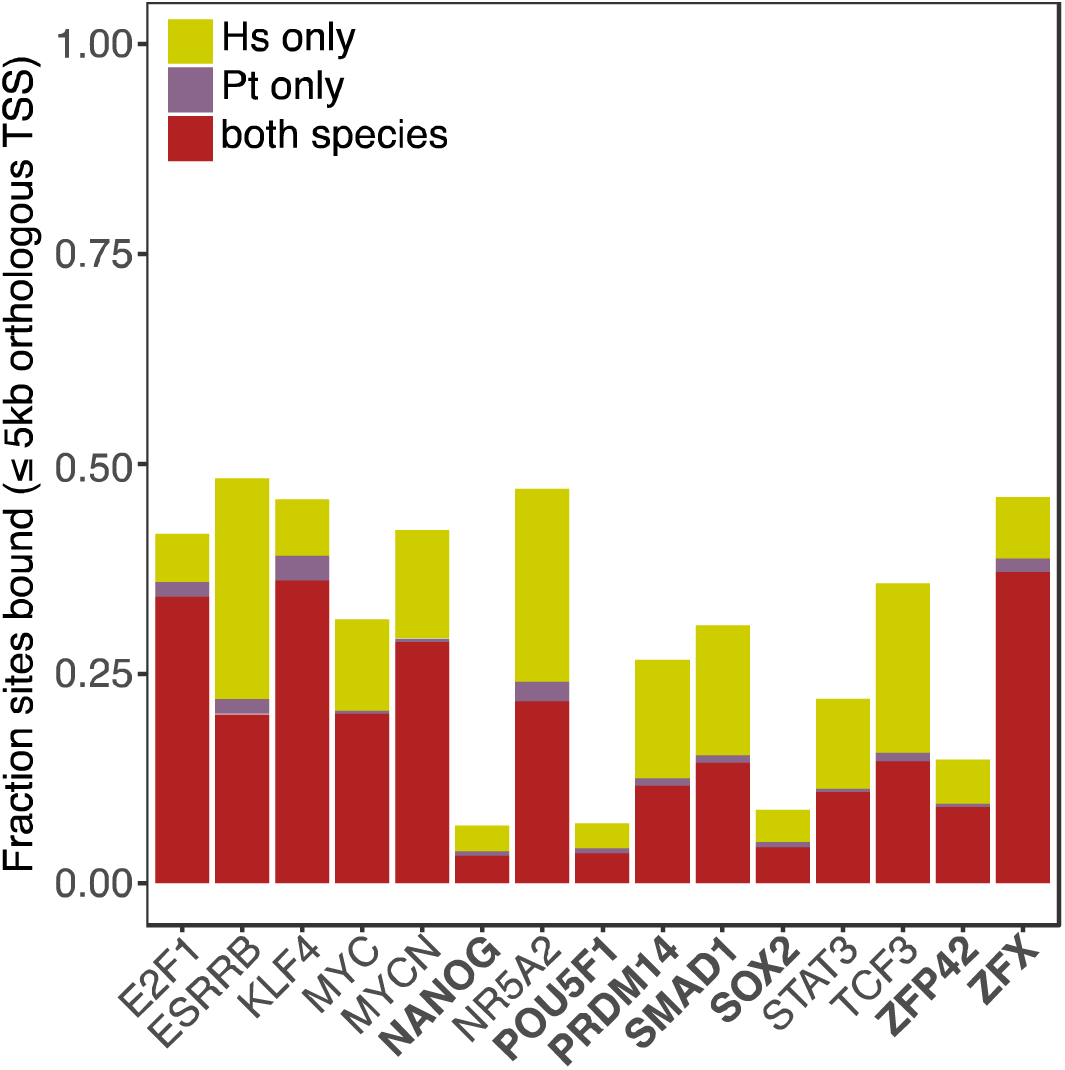
Predicted binding activity across 15 key pluripotency regulators genome-wide. Factors in boldface are associated with at least 50 sites with a PWM score ≥ 12.

## Discussion

The regulation of gene expression is a dynamic process orchestrated by multiple interacting cellular mechanisms. Transcription factors operate at the point of transcriptional initiation, by promoting (or sometimes inhibiting) the formation of the transcriptional complex through a suite of mechanisms. Here we have characterised and inferred the chromatin and TF binding landscapes in human and chimpanzee iPSCs. As might be expected given that we chose to use a cell type that mimics very early embryonic development, overall we observed high conservation of chromatin accessibility and TF binding in the two species. We found relatively few instances of interspecies regulatory differences that can be associated to downstream differences in expression. Taken together, our results consistently suggest that the pluripotency gene regulatory network is highly conserved between humans and chimpanzees – and seems highly robust to those interspecies differences we do observe. Indeed, there was little evidence to suggest that differences in chromatin architecture impact the pluripotency gene regulatory network that helps maintain cell identity. Much higher regulatory divergence is typically observed in differentiated tissues or cell types [21, 34] and indeed, we have also observed inter-species differences in chromatin accessibility near genes implicated in early developmental processes, such as embryonic patterning. The systematic difference in chromatin accessibility amongst bivalently modified genes known to be implicated in embryonic patterning is intriguing and, in our mind, worthy of further investigation.

We performed an in-depth examination of the potential of footprinting to infer genome-wide TF binding activity across scores of TFs simultaneously, using publicly available PWM datasets. There are variable levels of concordance between our binding predictions and ENCODE ChIP-seq data. While some of this discordance is attributable to technical artefacts, our data are in line with previous observations that PWMs do not always fully capture the binding behaviours of particular TFs [61]. In spite of this caveat, our results suggest that much of the binding activity is conserved between human and chimpanzees across the vast majority of PWMs we tested. This finding recapitulates a study of early development between the closely related *Drosophila melanogaster* and *Drosophila yakuba*, which found that these species shared between 85–98% of TFBS across six developmental regulators [62].

When considered at the scale of entire gene regulatory networks, we found strong evidence of redundancy in TF binding activity [36]. We recapitulated the observation that *cis*-acting turnover at possible binding sites is the main driver of divergence in binding activity [38], more than other possible forces such as *trans*-acting evolutionary divergence at the TF itself, or differences in TF expression levels. Indeed, the majority of the differences we identified, both in chromatin accessibility and TF binding, appear to be *cis*-acting. It has been proposed that *cis*-acting changes are more likely than *trans*-acting changes to drive evolutionary change, given the decreased likelihood that they will give rise to harmful pleiotropic effects, especially during development [2, 63].

We have previously characterised interspecies differences in human and chimpanzee iPSCs across a suite of regulatory mechanisms – DNA methylation, H3K27ac, H3K27me3 [44], transposable element activity [64] and now, chromatin accessibility and TF binding activity. Though it is difficult to compare divergence across different mechanisms given the multitude of approaches used to collect the data, it seems reasonably safe to state that divergence in all of the regulatory mechanisms we studied is consistently of lesser magnitude than the inter-species changes we observed in gene expression levels. It is unlikely that this is due to consistently worse resolution across this suite of assays than in RNA-seq. Even at the level of RNA-seq, we observed very little intra-specific variation in iPSCs relative to that seen in other somatic tissues, which lead to a dramatic increase in statistical power to identify differential gene expression between the two species [44]. Thus it might be the case that the pluripotent state is subject to much stronger evolutionary constraint than terminally differentiated downstream cell types.

## Methods

### Data collection and sequencing

We generated ATAC-seq libraries from 6 previously described chimpanzee iPSC lines and 6 previously described human iPSC lines [44]. All lines were cultured under feeder free conditions on hESC-grade Matrigel (Corning) and Essential 8 (Gibco) media as previously described. Paired end ATAC-seq libraries were generated as in [43] with a single exception: we collected 200,000 cells per pellet rather than the 50,000 described in the original protocol, solely because this made the pellet visible to the naked eye. Additionally, we collected two pellets at the same time for each cell line in case library preparations failed. Sample collection and library preparation were randomised with respect to species at all times. All libraries were multiplexed together and sequenced to an average depth of roughly 252 million reads across 3 flow cells of an Illumina HiSeq 2500; more details of the sequencing output are available in supplementary table 1.

### Read mapping, QC, and definition of high-confidence orthologous regions

To compare patterns of chromatin accessibility and TF binding between species we subjected all reads to a series of stringent QC and filtering steps. First, we mapped all reads independently to either the human (hg19) or chimpanzee (panTro 2.1.3) genomes using BWA 0.7.9a-r786 [65], allowing a maximum of two mismatches per read and a maximum fragment size of 5000 base pairs for paired-end mates. Reads with mapping quality < 30, unmapped reads, multi-mapping reads and reads that mapped outside the autosomes and X chromosome were discarded. We also discarded all reads that mapped to mitochondrial DNA for the main analyses reported in the text (but see below). Next, we identified and removed reads produced by PCR duplicates with Picard Tools (version 1.129; http://broadinstitute.github.io/picard/). The number of reads retained at each step is summarised in supplementary table 1.

Additionally, to control for differences in genome assembly quality, and to ensure fair comparisons between the two species, we retained only those reads that fell within a defined set of windows with high orthology between human and chimpanzee. The list of windows was generated by first retrieving the hg19toPanTro3 liftOver chain file from the UCSC Genome Browser [66], and using the regions that are part of the chain to generate non-overlapping 100 bp windows, for a total of 27.2 million 100bp windows. We then used liftOver [67] to test whether each of these windows could be lifted over to a single site in the chimpanzee genome, and from there back to its original location in the human genome, allowing for a maximum 20% change in size during each lift over process. Windows that failed either of these steps were discarded. In parallel, we calculated the uniqueness of every 50-mer in the chimpanzee and human genomes using the GEM suite [68]. Windows where greater than 20% of the 50-mers in either species were not uniquely mappable in their respective genomes, were removed from further analysis. In total, we retained ~17.6 million 100-bp high-quality windows for downstream analyses.

Each flow cell was processed separately at this stage, to allow us to identify any flow-cell or library-specific effects. Some libraries yielded very low numbers of mapped reads (< 100,000) and were clear outliers in QC. Although this is probably due to poor multiplexing rather than poor complexity, we discarded these libraries from all downstream analyses and replaced them with new libraries generated from the second pellet collected from the relevant individual. On the whole, we found overall robust reproducibility between libraries prepared from the same individual and thus present analyses at the individual level throughout. Ultimately, between 3.99 and 20.8% of generated raw reads per sample met all QC thresholds and were used in all of the following analyses, unless explicitly noted otherwise.

### Mitochondrial lineage reconstruction

During mapping we observed that a large fraction of reads (between 13.7% and 57.1%) were of mitochondrial origin. Because our chimpanzee iPSC lines are derived from second or third generation captive born individuals from the Yerkes Primate Research Centre, their genetic background is both admixed and unascertained. Therefore, we used MIRA 4 [69] and MITObim 1.8 [70] to assemble the mitochondrial genomes of all sequenced libraries. We made use of the publicly available pipeline at https://github.com/chrishah/MITObim, using the ‘quick’ option, and baited the assemblies with the human rCRS (Genbank: NC_012920) or the chimpanzee reference mtDNA genome (Genbank: NC_001643), as appropriate. We successfully generated single-contig mtDNA assemblies for all libraries. A maximum likelihood tree of the sequences, built with MEGA 7 [71], confirmed that in all cases except for one chimpanzee library, multiple libraries from the same individual grouped together as sister taxa (supplementary figure 5), confirming there were no sample swaps during sample preparation or sequencing. The one mismatched sample fell into a clade with a maternal cousin. In order to help establish the maternal lineage of the lines in our chimpanzee panel, we also included additional chimpanzee mtDNA sequences from [72] representing all four chimpanzee sub-species (supplementary figure 1).

### Identification of fragmentation biases

As crude indicators of quality of the ATAC-seq data, we examined the fragment size distribution of each library. Although libraries were prepared randomly with regards to species, there is a clear and consistent difference in fragmentation patterns between the two species, with chimpanzee libraries exhibiting an excess of short fragments (<100bp) originating in the nucleosome-free region relative to humans. It is unclear whether this reflects a biological or technical confounder, but to better quantify it we defined a series of *ad hoc* metrics and tallied the number of read pairs with fragment sizes between 50-59 bp, 100-109 bp, 150-159 bp and 190-199 bp, as well as the ratios between these measurements, and tested for associations between these measurements and all principal components in the data. In the orthologous TSS data, PC1 is strongly associated with the ratio of fragments 50-59 bp long to those 100-109 bp long (henceforth ratio 50/100, *P* = 1.7^∗^10^−4^, all values are available in supplementary table 2) and 50-59 bp to 150-159 bp (henceforth ratio 50/150, *P* = 3.4^∗^10^−5^), and PC2 is additionally associated with the ratio of reads mapping to orthologous TSS vs reads mapping to orthologous background regions (*P* = 3.7^∗^10^−4^), which may be an indicator of data noisiness. No PCs are associated with overall sequencing depth. When we consider the 5,675 ‘highly accessible’ regions the association with fragmentation bias, although diminished, remains significant (ratio 50/100 PC1 *P* = 4.2^∗^10^−4^, ratio 50/150 PC1 *P* = 0.04).

We note that cell lines were cultured together by the same individual (IGR), and randomised at all times relative to species to prevent the introduction of technical biases associated with our variable of interest. It is possible that this difference in fragmentation patterns is driven by increased sensitivity of chimpanzee iPSCs to the lysis buffers used in ATAC-seq, or by slightly different responses to our restrictive cell culture conditions. We have systematically elected to be conservative when interpreting our results, although in light of our thorough quality control pipeline we are confident that the trends we observe are reflective of true biological signal. Regardless of cause, this excess of short fragments in chimpanzees leads to a decrease in our ability to differentiate between open and closed chromatin in chimpanzees relative to humans, especially at small scales, and consequently to a loss of power to detect binding in chimpanzees relative to humans. What appears to be a human-unique event is likely to be actually conserved between the two species. For the same reason, those chimpanzee-unique accessibility events we do observe are likely to be true positives.

We considered subsampling the chimpanzee data to match the average human fragment distribution, but given both our uncertainty as to the cause of this difference and the relatively low number of reads that are retained given our exhaustive QC strategy, we reasoned that it was more likely to lead to an overall loss, rather than a gain, in power to detect differences.

### Characterising activity near transcription start sites

To examine patterns of activity near transcription start sites, we began by defining a set of highly orthologous meta-exons between human and chimpanzee as in [21], using Ensemblv75 (February 2014; code and documentation for meta-exon identification is available at [http://www.bitbucket.org/ee_reh_neh/orthoexon]; [73]). This data set contains at least 1 orthologous meta-exon associated with 40,075 human genes. We then defined the 5’-most position of the first meta-exon of each gene (adjusted for strand direction) as its orthologous transcription start site (orthologous TSS), and computed the number of high-quality orthologous 100bp windows (defined above) that fell within a 2kb window centred on the orthologous TSS. We retained only those regions with ≥ 50% overlap, which yielded a set of 28,238 orthologous TSSs. As an additional filtering step, we discarded 7,493 orthologous TSSs that did not span the annotated human TSS for the relevant gene according to Ensembl 75 GENCODE Basic [74] annotations (when multiple GENCODE Basic transcripts were associated with a single gene, we used the average location of all TSS). In total, we retained 20,745 orthologous TSSs.

In parallel, we defined a set of randomly selected 2kb orthologous background regions matched to the orthologous TSSs for broad mappability, additionally requiring that they be at least 5kb away from any annotated orthologous TSS. To identify regions with read depths suggestive of activity above the background cutting rate, we used k-means clustering to group the orthologous TSS data into three clusters on the basis of mean CPM by species. We set k = 3 rather than 2 to capture the long left tail of the distribution, which is readily visible in figure 1a.

### Differential expression and accessibility analyses

For consistency with our set of defined orthologous TSS we reanalysed previously published RNA-sequencing data from these cell lines using our updated set of orthologous exons. Analyses were performed using the same analysis script as previously [44]; overall we find 4,244 differentially expressed genes out of 12,674 testable genes, and high concordance between the old and new results (ρ= 0.89; past results included an additional human and chimpanzee individual). We intersected the expression data with our set of high-quality orthologous TSSs, and found a total of 5,675 genes with evidence for having an accessible orthologous TSS in at least one species – although the majority, 4,644, were accessible in both species. Of those regions seen only in one species, 873 are humans-unique, and 158 chimpanzee-unique. We tested the highly accessible regions for significant interspecies differences with limma [75] and the same steps that we used to test for differential expression. Given the partially confounded covariate structure, we considered four different models:

1. y ~ β_species_ + ε;
2. y ~ β_species_ + β_50/100_ + ε;
3. y ~ β_species_ + β_50/150_ + ε;
4. y ~ β_species_ + β_50/100_ + β_50/150_ + ε.

To identify the best model, we computed the AIC for each gene under each model, and found that models 1 and 3 were associated with the lowest AIC in roughly the same number of genes (1765 for model 1, 1909 for model 3), suggesting an overall greater suitability to the data. Given this result, we performed differential accessibility testing using model 3. However, we note that all four sets of results are qualitatively similar (supplementary figure 6).

### Estimating transcription factor binding with msCentipede

In order to determine a set of suitable PWM genomics matches to consider for analysis, we downloaded all 640 PWMs in version 10 of the HOCOMOCO database [54]. To ensure fair comparisons between human and chimpanzee data, we scanned both genomes for matches to each PWM, retaining any site with a PWM score >= 7. The adoption of this permissively low threshold was motivated by our desire to capture turnover at the PWM site between the two species. We then used liftOver [67] to ensure that identified human PWM matches were also present in the chimpanzee genome, and could be lifted over back to their original location in the human genome. We then used the k-mer mappability data described above to ensure that at least 80% of the 50-mers originating ± 100bp around the PWM site were uniquely mapping in both species. Finally, we excluded all PWMs associated with transcriptions factors that were either not present or not expressed in at least half of the individuals from one species according to our RNA-seq data, with the exception of master pluripotency regulators OCT4 and NANOG, where we confirmed expression through qPCR These two genes cannot be reliably assayed through RNA-seq due to the existence of closely related pseudogenes that confound mapping. We also included the *REX1* (also known as *ZFP42*) PWM from HOCOMOCO v11, as we had previously identified it as the sole master regulator of pluripotency that was differentially expressed between the two species. After all of these filtering steps, we retained 133,103,977 possible PWM sites across 306 TFs, 906,612 of which had a PWM score >= 12 in at least one species.

Finally, we used msCentipede [45], an extension of the CENTIPEDE algorithm [46] that can better capture heterogeneity around binding sites, to predict TF binding in all PWM sites in human and chimpanzee iPSCs. The vast majority of log posterior odds distributions reported by msCentipede are unimodal, with an extremely long rightward tail and without a clear break point (supplementary figure 7); chimpanzee distributions in particular are clearly narrower and left-shifted relative to human distributions, again reflecting our diminished power in chimpanzees. As such, we conservatively chose to call a site bound in a species if the log posterior odds were ≥ log (0.99/0.01) at that particular site, and unbound if they fell below that threshold. However, we note that our results are qualitatively robust to lowering this threshold to log (0.95/0.05) in chimpanzees to account for the decreased power in that species, resulting in an increase in the mean fraction of bound sites in chimpanzee, to 12.1%, a slight decrease in the fraction of sites called as bound in chimpanzee that are also bound in human, to 77.1%, and overall corroboration of our findings.

Additionally, HOCOMOCO summarises the quality and predictive value of PWMs using a simple A-B-C-D quality score, which is assigned to each PWM on the basis of receiver operating curve analyses [54]. Nearly half of the PWMs in the dataset (n = 133) have been assigned to class D, suggesting they might be information-poor and only capture a small subset of binding activity. This reflects broader observations that TFs can vary dramatically in their degree of preference for specific binding motifs [76]. We do find that the fraction of MPBS predicted to be bound does vary across PWM qualities, but only when we consider binding predictions in the ‘high-quality’ motif subset in chimpanzees (ANOVA *P* value = 0.016 genome-wide, 0.03 within 5kb of annotated orthologous TSS). In both of these cases, the significance is driven by differences between class A and D PWMs, but its overall impact appears relatively minor (supplementary figure 8). In light of these findings, we retained all PWMs for downstream analyses.

All analyses were performed using R 3.2.2 [77].

### Data accessibility

All raw ATAC-seq reads have been deposited in GEO/SRA under series number GSE122319, alongside individual msCentipede calls for all high quality sites and summary statistics for all tested PWMs. Previously published RNA-seq data from human and chimpanzee iPSCs is available under BioProject PRJNA260053. For comparisons with published ENCODE data we used all publicly released “optimal idr thresholded peaks” associated with either the human embryonic stem cell line H1 or the human induced pluripotent stem cell line GM23339 available on the ENCODE browser (https://www.encodeproject.org) as of the 15th of May, 2018. A full list of accession identifiers is provided in supplementary table 6.

## Acknowledgements

We thank members of the Gilad lab for helpful discussions, as well as Anil Raj and Heejung Shim for advice regarding msCentipede. IGR was supported in part by a Sir Henry Wellcome Postdoctoral Fellowship. SG was supported by Marie Skłodowska-Curie actions grant 655732-WhereWolf. This work was additionally supported by an NIGMS award to YG.

## Tables

Supplementary table 1: Sequencing depths and quality filtering steps.

Supplementary table 2: P-values for the association between each principal component and possible covariates

Supplementary table 3: Results of differential accessibility testing across 5,675 ‘highly accessible’ orthologous TSS using model 3 as described above.

Supplementary table 4: Results of differential expression testing across 12,674 orthologous genes.

Supplementary table 5: List of genes driving the GO enrichment results.

Supplementary table 6: Summary of ENCODE TF ChIP datasets used in this publication.

## Supplementary figures

Supplementary figure 1: Maximum likelihood tree from reconstructed and publicly available chimpanzee mtDNA sequences.

Supplementary figure 2: Principal component analysis of a. 20,745 orthologous transcription start sites genome-wide. b. 20,745 control regions genome-wide.

Supplementary figure 3: Log_2_ cpm distributions by individual around orthologous TSS and orthologous background regions.

Supplementary figure 4: Fraction of sites in the high-quality motif subset predicted to be bound within 5kb of an annotated orthologous TSS and genome wide.

Supplementary figure 5: Maximum likelihood tree from reconstructed human and chimpanzee mtDNA sequences.

Supplementary figure 6: overlap in differential accessibility results across the four considered models.

Supplementary figure 7: Log posterior binding probabilities for all 306 PWMs in the study, by species. The dashed vertical line indicates ln(0.99/0.01), our threshold for calling a site bound.

Supplementary figure 8: Fraction of sites predicted to be bound by PWM across the four main HOCOMOCO quality classes. Pairwise test P-values are reported after Tukey’s post hoc HSD correction.

## Abbreviations

ATAC: assay for transposase-accessible chromatin
CPM: counts per million
DA: differentially accessible
DE: differentially expressed
ESC: embryonic stem cell
FDR: false discovery rate
iPSC: induced pluripotent stem cell
MPBS: motif-predicted binding site(s)
PCA: principal component analysis
PWM: position weight matrix
RPKM: reads per kilobase per million reads mapped
TF: transcription factor
TFBS: transcription factor binding site(s)
TSS: transcription start site(s)

## References

1. King, M. and A. Wilson, Evolution at two levels in humans and chimpanzees. Science, 1975. 188(4184): p. 107-116.

2. Carroll, S.B., Evolution at two levels: On genes and form. PLoS Biology, 2005. 3(7): p. 1159-1166.

3. Britten, R.J. and E.H. Davidson, Repetitive and non-repetitive DNA sequences and a speculation on the origins of evolutionary novelty. Quarterly Review of Biology, 1971. 46(2): p. 111-138.

4. Jacob, F., Evolution and tinkering. Science, 1977. 196(4295): p. 1161-6.

5. Gallego Romero, I., I. Ruvinsky, and Y. Gilad, Comparative studies of gene expression and the evolution of gene regulation. Nature Reviews Genetics, 2012. 13(7): p. 505-516.

6. Bejerano, G., et al., Ultraconserved elements in the human genome. Science, 2004. 304(5675): p. 1321-5.

7. Siepel, A., et al., Evolutionarily conserved elements in vertebrate, insect, worm, and yeast genomes. Genome Research, 2005. 15(8): p. 1034-50.

8. Prabhakar, S., et al., Accelerated evolution of conserved noncoding sequences in humans. Science, 2006. 314(5800): p. 786.

9. Pollard, K.S., et al., Forces shaping the fastest evolving regions in the human genome. PLoS Genetics, 2006. 2(10): p. e168.

10. Booker, B.M., et al., Bat Accelerated Regions Identify a Bat Forelimb Specific Enhancer in the HoxD Locus. PLoS Genetics, 2016. 12(3): p. e1005738-e1005738.

11. Eckalbar, W.L., et al., Transcriptomic and epigenomic characterization of the developing bat wing. Nature Genetics, 2016. 48(5): p. 528-536.

12. Pollard, K., et al., An RNA gene expressed during cortical development evolved rapidly in humans. Nature, 2006. 443(7108): p. 167-172.

13. Hubisz, M.J. and K.S. Pollard, Exploring the genesis and functions of Human Accelerated Regions sheds light on their role in human evolution. Current Opinion in Genetics and Development, 2014. 29: p. 15-21.

14. Kvon, E.Z., et al., Progressive Loss of Function in a Limb Enhancer during Snake Evolution. Cell, 2016. 167(3): p. 633-642.e11.

15. O’Brown, N.M., et al., A recurrent regulatory change underlying altered expression and Wnt response of the stickleback armor plates gene EDA. Elife, 2015. 4: p. e05290.

16. Reilly, S., et al., Evolutionary changes in promoter and enhancer activity during human corticogenesis. Science, 2015. 347(6226): p. 1155-1159.

17. Bakken, T.E., et al., A comprehensive transcriptional map of primate brain development. Nature, 2016. 535(7612): p. 367-375.

18. Cotney, J., et al., The Evolution of Lineage-Specific Regulatory Activities in the Human Embryonic Limb. Cell, 2013. 154(1): p. 185-196.

19. Prabhakar, S., et al., Human-Specific Gain of Function in a Developmental Enhancer. Science, 2008. 321(5894): p. 1346-1350.

20. Blekhman, R., et al., Gene regulation in primates evolves under tissue-specific selection pressures. PLoS Genetics, 2008. 4(11): p. e1000271-e1000271.

21. Blekhman, R., et al., Sex-specific and lineage-specific alternative splicing in primates. Genome Research, 2010. 20(2): p. 180-189.

22. Brawand, D., et al., The evolution of gene expression levels in mammalian organs. Nature, 2011. 478(7369): p. 343-348.

23. Khaitovich, P., et al., Regional patterns of gene expression in human and chimpanzee brains. Genome Research, 2004. 14(8): p. 1462-1473.

24. Enard, W., et al., Intra- and interspecific variation in primate gene expression patterns. Science, 2002. 296(5566): p. 340-3.

25. He, Z., et al., Comprehensive transcriptome analysis of neocortical layers in humans, chimpanzees and macaques. Nature Neuroscience, 2017.

26. Cain, C.E., et al., Gene expression differences among primates are associated with changes in a histone epigenetic modification. Genetics, 2011. 187(4): p. 1225-1234.

27. Zhou, X., et al., Epigenetic modifications are associated with inter-species gene expression variation in primates. Genome Biology, 2014. 15(12): p. 547-547.

28. Pai, A., et al., A Genome-Wide Study of DNA Methylation Patterns and Gene Expression Levels in Multiple Human and Chimpanzee Tissues. PLOS Genetics, 2011. 7(2): p. e1001316-e1001316.

29. Hernando-Herraez, I., et al., The interplay between DNA methylation and sequence divergence in recent human evolution. bioRxiv, 2015: p. 015966-015966.

30. Hernando-Herraez, I., et al., Dynamics of DNA Methylation in Recent Human and Great Ape Evolution. PLoS Genetics, 2013. 9(9): p. e1003763-e1003763.

31. Bilgin Sonay, T., et al., Tandem repeat variation in human and great ape populations and its impact on gene expression divergence. Genome Research, 2015. 25(11): p. 1591-9.

32. Blekhman, R., A. Oshlack, and Y. Gilad, Segmental duplications contribute to gene expression differences between humans and chimpanzees. Genetics, 2009. 182(2): p. 627-630.

33. Ward, M.C., et al., Silencing of transposable elements may not be a major driver of regulatory evolution in primate iPSCs. Elife, 2018. 7.

34. Nowick, K., et al., Differences in human and chimpanzee gene expression patterns define an evolving network of transcription factors in brain. Proc Natl Acad Sci U S A, 2009. 106(52): p. 22358-63.

35. Shibata, Y., et al., Extensive Evolutionary Changes in Regulatory Element Activity during Human Origins Are Associated with Altered Gene Expression and Positive Selection. PLOS Genetics, 2012. 8(6): p. e1002789-e1002789.

36. Berthelot, C., et al., Complexity and conservation of regulatory landscapes underlie evolutionary resilience of mammalian gene expression. Nat Ecol Evol, 2018. 2(1): p. 152-163.

37. Paris, M., et al., Extensive divergence of transcription factor binding in Drosophila embryos with highly conserved gene expression. PLoS Genet, 2013. 9(9): p. e1003748.

38. Schmidt, D., et al., Five-vertebrate ChIP-seq reveals the evolutionary dynamics of transcription factor binding. Science, 2010. 328(5981): p. 1036-40.

39. Wilson, M.D., et al., Species-specific transcription in mice carrying human chromosome 21. Science, 2008. 322(5900): p. 434-8.

40. Li, X.Y., et al., The role of chromatin accessibility in directing the widespread, overlapping patterns of Drosophila transcription factor binding. Genome Biol, 2011. 12(4): p. R34.

41. Peng, P.-C., et al., Evolutionary changes in DNA accessibility and sequence predict divergence of transcription factor binding and enhancer activity. bioRxiv, 2018.

42. Vierstra, J. and J.A. Stamatoyannopoulos, Genomic footprinting. Nature Methods, 2016. 13(3): p. 213-221.

43. Buenrostro, J.D., et al., Transposition of native chromatin for fast and sensitive epigenomic profiling of open chromatin, DNA-binding proteins and nucleosome position. Nature Methods, 2013. 10(12): p. 1213-1218.

44. Gallego Romero, I., et al., A panel of induced pluripotent stem cells from chimpanzees: a resource for comparative functional genomics. eLife, 2015. 4: p. e07103-e07103.

45. Raj, A., et al., msCentipede: Modeling Heterogeneity across Genomic Sites and Replicates Improves Accuracy in the Inference of Transcription Factor Binding. PLOS ONE, 2015. 10(9): p. e0138030-e0138030.

46. Pique-Regi, R., et al., Accurate inference of transcription factor binding from DNA sequence and chromatin accessibility data. Genome Research, 2011. 21(3): p. 447-455.

47. Ashburner, M., et al., Gene Ontology: tool for the unification of biology. Nature Genetics, 2000. 25(1): p. 25-29.

48. Rada-Iglesias, A., et al., A unique chromatin signature uncovers early developmental enhancers in humans. Nature, 2010. 470(7333): p. 279-283.

49. Roadmap Epigenomics, C., et al., Integrative analysis of 111 reference human epigenomes. Nature, 2015. 518(7539): p. 317-30.

50. Bu, L., et al., Human ISL1 heart progenitors generate diverse multipotent cardiovascular cell lineages. Nature, 2009. 460(7251): p. 113-7.

51. McCormick, M.B., et al., NeuroD2 and neuroD3: distinct expression patterns and transcriptional activation potentials within the neuroD gene family. Mol Cell Biol, 1996. 16(10): p. 5792-800.

52. Masserdotti, G., et al., Transcriptional Mechanisms of Proneural Factors and REST in Regulating Neuronal Reprogramming of Astrocytes. Cell Stem Cell, 2015. 17(1): p. 74-88.

53. Tsuda, H., et al., Structure and promoter analysis of Math3 gene, a mouse homolog of Drosophila proneural gene atonal. Neural-specific expression by dual promoter elements. J Biol Chem, 1998. 273(11): p. 6327-33.

54. Kulakovskiy, I.V., et al., HOCOMOCO: Expansion and enhancement of the collection of transcription factor binding sites models. Nucleic Acids Research, 2016. 44(D1): p. D116-D125.

55. Consortium, E.P., An integrated encyclopedia of DNA elements in the human genome. Nature, 2012. 489(7414): p. 57-74.

56. Young, R.A., Control of the Embryonic Stem Cell State. Cell, 2011. 144(6): p. 940-954.

57. Marchetto, M.C.N., et al., Differential L1 regulation in pluripotent stem cells of humans and apes. Nature, 2013. 503(7477): p. 525-529.

58. Chronis, C., et al., Cooperative Binding of Transcription Factors Orchestrates Reprogramming. Cell, 2017. 168(3): p. 442-459 e20.

59. Orkin, S.H. and K. Hochedlinger, Chromatin Connections to Pluripotency and Cellular Reprogramming. Cell, 2011. 145(6): p. 835-850.

60. Ng, H.-H. and M. Surani, The transcriptional and signalling networks of pluripotency. Nat Cell Biol, 2011. 13(5): p. 490-496.

61. Weirauch, M.T., et al., Determination and inference of eukaryotic transcription factor sequence specificity. Cell, 2014. 158(6): p. 1431-1443.

62. Bradley, R.K., et al., Binding site turnover produces pervasive quantitative changes in transcription factor binding between closely related Drosophila species. PLoS Biol, 2010. 8(3): p. e1000343.

63. Wray, G., The evolutionary significance of cis-regulatory mutations. Nature Reviews Genetics, 2007. 8(3): p. 206-216.

64. Ward, M.C., et al., Silencing of transposable elements may not be a major driver of regulatory evolution in primate induced pluripotent stem cells. bioRxiv, 2017: p. 1-46.

65. Li, H. and R. Durbin, Fast and accurate short read alignment with Burrows-Wheeler transform. Bioinformatics, 2009. 25(14): p. 1754-1760.

66. Kent, W.J., et al., The human genome browser at UCSC. Genome Res, 2002. 12(6): p. 996-1006.

67. Hinrichs, A.S., et al., The UCSC Genome Browser Database: update 2006. Nucleic Acids Res, 2006. 34(Database issue): p. D590-8.

68. Derrien, T., et al., Fast computation and applications of genome mappability. PLoS ONE, 2012. 7(1): p. e30377-e30377.

69. Chevreux, B., T. Wetter, and S. Suhai, Genome Sequence Assembly Using Trace Signals and Additional Sequence Information, in Computer Science and Biology: Proceedings of the German Conference on Bioinformatics. 1999. p. 45-56.

70. Hahn, C., L. Bachmann, and B. Chevreux, Reconstructing mitochondrial genomes directly from genomic next-generation sequencing reads - A baiting and iterative mapping approach. Nucleic Acids Research, 2013. 41(13): p. e129--e129.

71. Tamura, K., et al., MEGA5: molecular evolutionary genetics analysis using maximum likelihood, evolutionary distance, and maximum parsimony methods. Mol Biol Evol, 2011. 28(10): p. 2731-9.

72. Bjork, A., et al., Evolutionary history of chimpanzees inferred from complete mitochondrial genomes. Molecular Biology and Evolution, 2011. 28(1): p. 615-623.

73. Yates, A., et al., Ensembl 2016. Nucleic Acids Res, 2016. 44(D1): p. D710-6.

74. Harrow, J., et al., GENCODE: the reference human genome annotation for The ENCODE Project. Genome Res, 2012. 22(9): p. 1760-74.

75. Ritchie, M., et al., limma powers differential expression analyses for RNA-sequencing and microarray studies. Nucleic Acids Research, 2015. 43(7): p. e47--e47.

76. Karimzadeh, M. and M.M. Hoffman, Virtual ChIP-seq: predicting transcription factor binding by learning from the transcriptome. bioRxiv, 2018.

77. R Core Team, R: A Language and Environment for Statistical Computing, R Foundation for Statistical Computing, Editor. 2015: Vienna.

